# Kinfitr – an open source tool for reproducible PET modelling: validation and evaluation of test-retest reliability

**DOI:** 10.1101/2020.02.20.957738

**Authors:** Jonathan Tjerkaski, Simon Cervenka, Lars Farde, Granville James Matheson

## Abstract

In positron emission tomography (PET) imaging, binding is typically estimated by fitting pharmacokinetic models to the series of measurements of radioactivity in the target tissue following intravenous injection of a radioligand. However, there are multiple different models to choose from and numerous analytical decisions which must be made when modelling PET data. Therefore, full communication of all the steps involved is often not feasible within the confines of a scientific publication. As such, there is a need to improve analytical transparency. *Kinfitr*, written in the open-source programming language R, is a tool developed for flexible and reproducible kinetic modelling of PET data, i.e. performing all steps using code which can be publicly shared in analysis notebooks. In this study, we compared outcomes obtained using *kinfitr* with those obtained using PMOD: a widely-used commercial tool.

Using previously-collected test-retest data obtained with four different radioligands, a total of six different kinetic models were fitted to time-activity curves derived from different brain regions. We observed high agreement between the two kinetic modelling tools both for binding estimates and for microparameters. Likewise, no substantial differences were observed in the test-retest reliability estimates between the two tools.

In summary, we showed excellent agreement between the open source R package *kinfitr*, and the widely-used commercial application PMOD. We therefore conclude that *kinfitr* is a valid and reliable tool for kinetic modelling of PET data.

## Background

Positron emission tomography (PET) is an imaging modality with high sensitivity and specificity for biochemical markers and metabolic processes *in vivo* [1]. It is an important tool in the study of psychiatric and neurological diseases, as well as for evaluating novel and established pharmacological treatments [2–4]. In PET imaging, study participants receive an intravenous injection of a radioligand, which binds specifically to a target molecule [5]. The concentration of radioligand in a region of interest (ROI) is measured over time to produce a time-activity curve (TAC) [6]. Radioligand binding, and thereby the concentration of the target molecule, can then be estimated using quantitative kinetic models [7, 8], of which there are many.

Importantly, the choice of a certain kinetic modelling approach should be based on several considerations, including the pharmacokinetic properties of the radioligand, the signal-to-noise ratio of the TAC, the availability of arterial blood sampling and the biological research question. Furthermore, there are various other analytical decisions that must be made in conjunction with modelling, such as the selection of statistical weighting. The sheer number of options available for kinetic modelling, in addition to those in prior pre-processing of image data [9] and blood data [10, 11] means that the communication of all analytical steps may not be feasible within the confines of a scientific publication. This limitation may, in turn, impede replication efforts and obscure potential errors [12]. Such problems have been described in numerous fields, and reproducible research practices have been proposed as a solution: this means increasing transparency by exposing more of the research workflow to the scientific community, through sharing of code and (when possible) data [12–14].

Several tools, both commercial and open-source, have been developed to facilitate the analysis of PET data [15–18]. These tools differ in their focus on various levels of analysis such as image reconstruction, image processing or high-throughput quantification. *Kinfitr* is an open source software package specifically developed for the purpose of performing PET kinetic modelling in a flexible and reproducible fashion. It is written in the R programming language [19], which provides access to a rich ecosystem of tools for reproducible research. The overall aims of *kinfitr* are to provide researchers with a high degree of flexibility during modelling as well as to provide the user with the ability to report all the steps taken during this process in a transparent manner [20]. This software package has been used in several scientific publications [21–24], however, it has not yet been formally evaluated against other software. This is an important safeguard for open-source software, as bugs can otherwise go unnoticed (e.g. [25]).

The purpose of this study was to validate *kinfitr* by comparing its estimates to those obtained with the widely-used commercially available software PMOD [17], which for the purposes of this analysis was considered to be the gold standard within the field. Making use of previously collected test-retest data for four different radioligands, we evaluated the agreement between these tools, using three different kinetic models each.

## Methods

### Data and study participants

This study was performed using data from four previous studies carried out at the Centre for Psychiatry Research, Department of Clinical Neuroscience, Karolinska Institutet, Stockholm, Sweden. In all studies, the data collection was approved by the Regional Ethics and Radiation Safety Committee of the Karolinska Hospital, and all subjects had provided written informed consent prior to their participation. All participants were young (aged 20-35 years), healthy individuals who underwent two PET measurements each with the same radioligand. The radioligands used were [^11^C]SCH23390 [26], [^11^C]AZ10419369 [27], [^11^C]PBR28 [28] and (R)- [^11^C]PK11195 [29]. Data from two target ROIs were selected as representative for each dataset. The two ROIs correspond to a region with higher and a region with lower specific binding for the radioligand used.

The [^11^C]SCH23390 cohort consisted of fifteen male subjects [30]. [^11^C]SCH23390 binds to the dopamine D1 receptor, which is highly concentrated in the striatum, with a lower concentration in cortical regions and negligible expression in the cerebellum [31]. In this study, the target ROIs were the striatum and the frontal cortex.

The [^11^C]AZ10419369 cohort consisted of eight male subjects [32]. [^11^C]AZ10419369 binds to the serotonin 5-HT_1B_ receptor, which is highly concentrated in the occipital cortex, with a moderate concentration in the frontal cortex and negligible expression in the cerebellum. The occipital and frontal cortices were selected as the target ROIs for [^11^C]AZ10419369 [32].

The [^11^C]PBR28 cohort consisted of 6 males and 6 females[33] and the (R)-[^11^C]PK11195 cohort was comprised of 6 male individuals[34]. Both [^11^C]PBR28 and (R)- [^11^C]PK11195 bind to the 18 kDa translocator protein (TSPO), a proposed marker of glial cell activation [35–37]. TSPO has a widespread distribution across the whole brain, predominantly in grey matter [38]. In this study, the ROIs used for both TSPO ligands were the thalamus and the frontal cortex. Furthermore, arterial blood sampling, plasma measurements and plasma metabolite analysis were performed and used in the analysis for the [^11^C]PBR28 and (R)- [^11^C]PK11195 cohorts as described previously [33, 34], as no true reference region is available for these radioligands.

### Kinetic modelling

In total, a total of six commonly-used kinetic models were used to quantify radioligand binding in the different datasets. For each analysis, both *kinfitr* (version 0.4.3) and PMOD (version 3.704, PMOD Technologies LLC., Zürich, Switzerland) was used. These estimates were subsequently compared to assess the agreement between the two kinetic modelling tools. The same investigator (JT) performed the analysis with both tools.

For the quantification of [^11^C]SCH23390 and [^11^C]AZ10419369, the Simplified Reference Tissue Model (SRTM) [39], Ichise’s Multilinear Reference Tissue Model 2 (MRTM2) [40] and the non-invasive Logan plot [41] were used, with the cerebellum as a reference region for both radioligands. These models will be referred to as the “reference tissue models”, whose main outcome was the binding potential (BP_ND_). Prior to performing MRTM2, k_2_’ was estimated by fitting MRTM1 [40] for the TAC of the higher-binding region for each subject, the result of which was used as an input when fitting MRTM2 for all regions of that particular subject.

For the quantification of (R)- [^11^C]PK11195 and [^11^C]PBR28, the two-tissue compartment model (2TCM) [42–44], the Logan plot [45] and Ichise’s Multilinear Analysis 1 (MA1) [46] were used to estimate the distribution volume (V_T_) using the metabolite-corrected arterial plasma (AIF) as an input function. These will henceforth be referred to as the “invasive models”. The delay between the TACs and arterial input function was fitted by the 2TCM using the TAC for the whole brain ROI. The default values in PMOD for the blood volume fraction (v_B_) were maintained throughout all analyses, which amounted to a v_B_ = 0 for MA1 and the invasive Logan plot and v_B_ = 0.05 for 2TCM.

The manner by which the analysis was performed was based on the explicit instructions provided along with each tool. However, when no explicit instructions were available, we resorted to making inferences based on the instructions for previous analytical steps and the design of the user interface of each kinetic modelling tool to emulate best how users might actually use each tool. For instance, one difference between how both tools are used relates to the selection of t*, which is required when fitting the linearized models (MA1, MRTM2 and both invasive and non-invasive Logan plots). These linearized models rely on asymptotic approximations, and t* is the time point after which these approximations apply. In *kinfitr*, a single t* value was selected and used across individuals, while in PMOD, a unique t* value was selected for each individual PET measurement. In both cases, the design of the software makes it more difficult and time-consuming to do this the other way (more details provided in Supplementary Materials S1), and in the former case this was a deliberate design decision to prevent over-fitting [20]. Importantly, the decision to focus on how the tools might be used in practice, rather than simply optimising the similarity of processing, provides more information about the extent to which outcomes might differ between tools, rather than the extent to which they might be made to be the same. We believe that this is of greater relevance to the research community.

### Statistics

The primary aim of this study was to assess the agreement between estimates of BP_ND_ (for reference tissue models) or V_T_ (for invasive models) obtained using *kinfitr* or PMOD, using a total of 6 different kinetic models. By using test-retest data, we were also able to evaluate the secondary aim of comparing the test-retest reliability within individuals for each tool.

The agreement between *kinfitr* and PMOD was evaluated using the intraclass correlation coefficient (ICC), the Pearson correlation coefficient, and bias.

The ICC represents the proportion of the total variance which is not attributable to measurement error, or noise. Therefore, an ICC of 1 represents perfect agreement, while an ICC of 0 represents no signal and only noise. We used the ICC(A,1) [47], which is computed using the following equation:

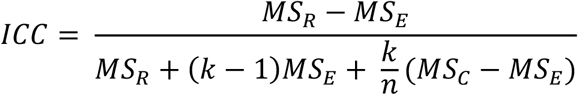

where MS_R_ is the mean sum of squares of the rows, MS_*E*_ is the mean sum of squares of the error and MS_C_ is the mean sum of squares of the columns; and where k refers to the number of raters or observations per subject (in this case 2), and n refers to the number of subjects [48].

Bias was defined as the percentage change in the means of the values of the binding estimates. This measure was calculated as follows:

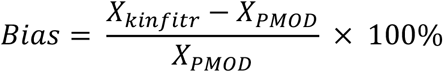

where X represents estimates of radioligand binding.

To compare the performance of each tool for assessing within- and between-subject variability, we calculated the mean, coefficient of variation (CV), ICC, within-subject coefficient of variation (WSCV) and absolute variability (AV).

The CV is calculated as a measure of dispersion. It is defined as follows:

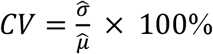

Where 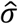 represents the sample standard deviation and 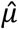 the sample mean of the binding estimate value.

The ICC was calculated as above, since inter-rater agreement and test-retest reliability are both most appropriately estimated using the two-way mixed effects, absolute agreement, single rater/measurement ICC, the ICC(A,1) [49].

The within-subject coefficient of variation was calculated as a measure of repeatability and expresses the error as a percentage of the mean. It is calculated as follows:

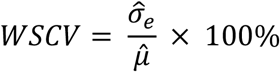

where 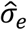 represents the standard error of the binding estimate value, which is analogous to the square root of the within subject mean sum of squares (MS_W_), which is also used in the calculation of the ICC above. 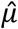 is the sample mean of the binding estimate value.

Finally, we also calculated the absolute variability (AV). This metric can be considered as an approximation of the WSCV above. While not as useful as the WSCV [50], AV has traditionally been applied within the PET field, and is included for historical comparability.

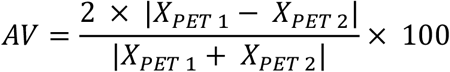

Where “X” refers to the value of the binding estimate and “PET 1” and “PET 2” refer to the first and second PET measurements in a test-retest experiment (in chronological order).

### Exclusions and Deviations

All subjects in the [^11^C]SCH23390, [^11^C]AZ10419369 and (R)- [^11^C]PK11195 cohorts were included in the final analysis. However, one study participant belonging to the [^11^C]PBR28 cohort, was excluded due to exhibiting a poor fit in the PMOD analysis which resulted in an abnormally high V_T_ estimate (>5 standard deviations from the mean of the rest of the sample, and a >500% increase from the other measurement of the same individual) (Supplementary Materials S2). We were unable to resolve this problem using different starting, upper and lower limits.

Moreover, in the analysis of the [^11^C]PBR28 cohort, *kinfitr* returned warnings about high values of k_3_ and k_4_ for some fits. When parameter estimates are equal to upper or lower limit bounds, the software recommends either altering the bounds, or attempting to use multiple starting points to increase the chance of finding the global minimum as opposed to a local minimum. Since in this case we deemed the values to be abnormally high, we opted for the latter strategy using the multiple starting point functionality of *kinfitr* using the *nls.multstart* package [51]. This entails fitting each curve a given number of times (we selected 100) using randomly sampled starting parameters from across the parameter space. This process led to negligible changes in the V_T_ estimates, but yielded microparameter estimates whose values were no longer equal to the upper or lower limit bounds.

### Data and Code Availability

All analysis code is available at https://github.com/tjerkaskij/agreement_kinfitr_pmod. The data are pseudonymized according to national (Swedish) and EU legislation and cannot be fully anonymized, and therefore cannot be shared openly within this repository due to current institutional restrictions. Metadata can be openly published, and the underlying data can instead be made available upon request on a case by case basis as allowed by the legislation and ethical permits. Requests for access can be made to the Karolinska Institutet’s Research Data Office at.

## Results

We found excellent agreement between binding estimates computed using both tools, with a median ICC of 0.98 (range: 0.81-1.00) (Table 1, Supplementary Materials S3) [50]. Likewise, we found high correlations between *kinfitr* and PMOD, with a median correlation coefficient of 0.99 (range: 0.95-1.00) (Table 1). It was observed that the linearized methods (i.e. MA1, MRTM2 and both invasive and non-invasive Logan plots) generally exhibited lower agreement than the non-linear models.

**Table 1:**
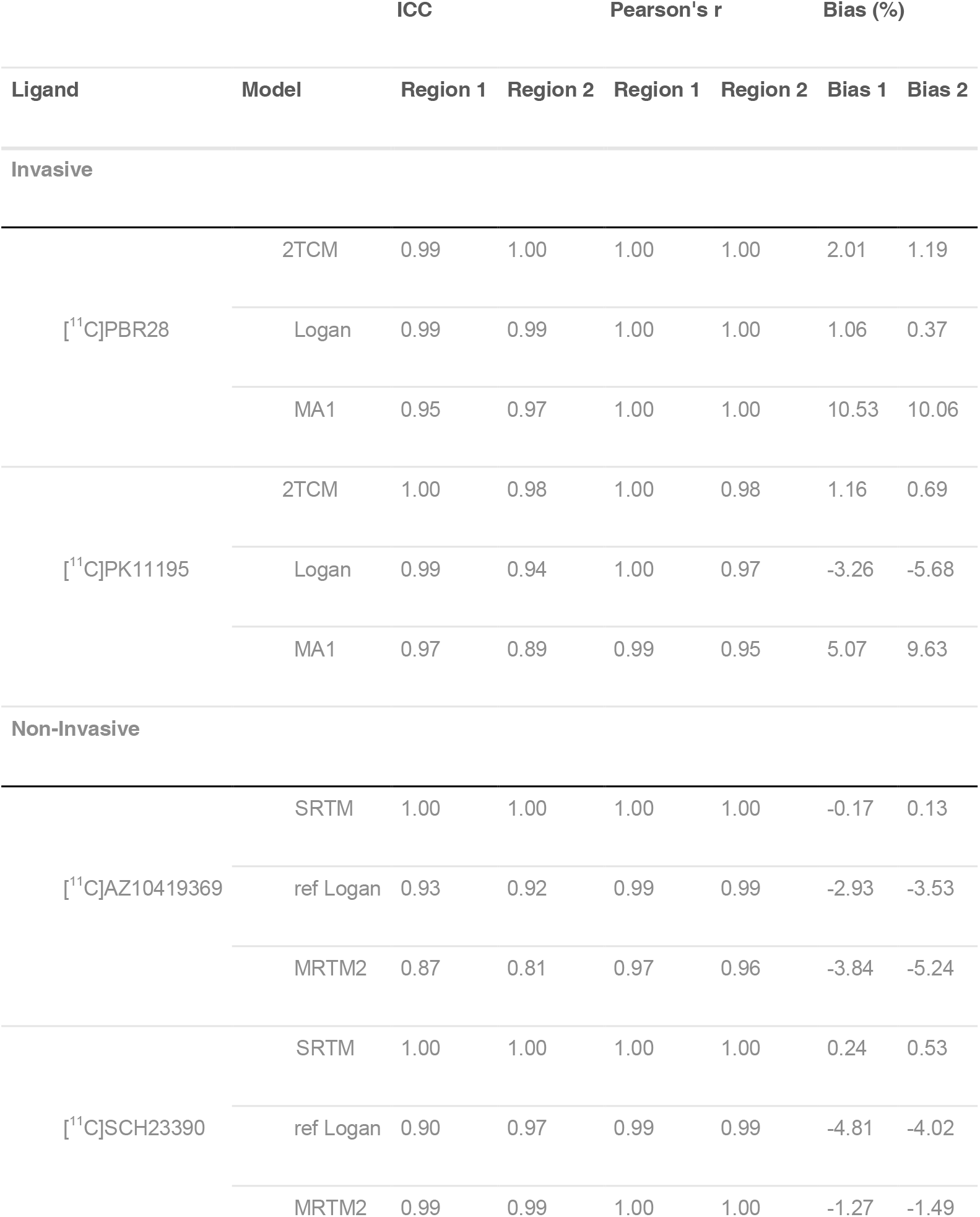
Agreement between *kinfitr* and PMOD. Region 1 corresponds to the occipital cortex for the radioligand [^11^C]AZ10419369, the striatum for [^11^C]SCH23390 and the thalamus for both (R)- [^11^C]PK11195 and [^11^C]PBR28. Region 2 corresponds to the frontal cortex for all four radioligands which were used in this study. Abbreviations: “2TCM” = Two-tissue compartmental model, “Logan” = Invasive Logan plot, “MA1” = Ichise’s Multilinear Analysis 1, “SRTM” = simplified reference tissue model, “ref Logan” = reference tissue Logan plot, “MRTM2” = Ichise’s Multilinear Reference Tissue Model 2 (MRTM2), “ICC” = intra-class correlation coefficient, “Pearson’s r” = Pearson’s correlation coefficient.

We also found strong correlations between the binding estimates of the different kinetic models that were estimated using *kinfitr* and PMOD (Supplementary Materials S4). When comparing the binding estimates of the three reference tissue models within *kinfitr* and PMOD respectively, there was a median Pearson’s correlation coefficient of 0.99 for both tools. For the invasive models, there was a median Pearson’s correlation coefficient of 0.79 for PMOD and 0.99 for *kinfitr*.

### Test-retest reliability

In general, both tools performed similarly, with no substantial differences seen in the mean values, dispersion (CV), reliability (ICC), or variability (WSCV and AV) (Table 2; Figure 1).

**Table 2:** Assessment of test-retest reliability of kinfitr and PMOD for a single high-binding ROI for each radioligand. The occipital cortex region was used for the radioligand [^11^C]AZ10419369, the striatum for [^11^C]SCH23390 and the thalamus for both (R)- [^11^C]PK11195 and [^11^C]PBR28. Abbreviations: “2TCM” = Two-tissue compartmental model, “Logan” = Invasive Logan plot, “MA1” = Ichise’s Multilinear Analysis 1, “SRTM” = simplified reference tissue model, “ref Logan” = reference tissue Logan plot, “MRTM2” = Ichise’s Multilinear Reference Tissue Model 2 (MRTM2), “ICC” = intra-class correlation coefficient, “CV” = Coefficient of variance, “WSCV” = within-subject coefficient of variance, “AV” = absolute variability.

| Ligand | Software | Model | Mean | CV (%) | ICC | WSCV (%) | AV (%) |
| --- | --- | --- | --- | --- | --- | --- | --- |
| Invasive |  |  |  |  |  |  |  |
| <sup>[11]C</sup>PK11195 | *kinfitr* | 2TCM | 0.76 | 27.1 | 0.75 | 13.9 | 18.9 |
|  | PMOD | 2TCM | 0.75 | 27.4 | 0.72 | 15.0 | 20.0 |
|  | *kinfitr* | Logan | 0.76 | 27.2 | 0.79 | 12.9 | 17.2 |
|  | PMOD | Logan | 0.79 | 26.2 | 0.80 | 12.2 | 16.2 |
|  | *kinfitr* | MA1 | 0.84 | 26.4 | 0.73 | 14.1 | 17.9 |
|  | PMOD | MA1 | 0.80 | 28.2 | 0.67 | 16.6 | 19.2 |
| <sup>[11]C</sup>PBR28 | *kinfitr* | 2TCM | 3.84 | 59.6 | 0.91 | 18.4 | 25.1 |
|  | PMOD | 2TCM | 3.73 | 57.3 | 0.89 | 19.1 | 26.7 |
|  | *kinfitr* | Logan | 3.75 | 57.4 | 0.91 | 17.7 | 24.4 |
|  | PMOD | Logan | 3.68 | 54.8 | 0.88 | 19.4 | 26.5 |
|  | *kinfitr* | MA1 | 4.00 | 58.4 | 0.91 | 18.0 | 24.0 |
|  | PMOD | MA1 | 3.54 | 53.1 | 0.91 | 16.7 | 23.7 |
| Non-Invasive |  |  |  |  |  |  |  |
| <sup>[11]C</sup>AZ10419369 | *kinfitr* | SRTM | 1.59 | 10.7 | 0.67 | 6.3 | 5.9 |
|  | PMOD | SRTM | 1.60 | 11.0 | 0.67 | 6.5 | 5.9 |
|  | *kinfitr* | ref Logan | 1.45 | 8.1 | 0.61 | 5.2 | 4.8 |
|  | PMOD | ref Logan | 1.49 | 8.3 | 0.62 | 5.3 | 5.5 |
|  | *kinfitr* | MRTM2 | 1.42 | 8.2 | 0.52 | 5.9 | 4.7 |
|  | PMOD | MRTM2 | 1.48 | 8.0 | 0.59 | 5.2 | 5.5 |
| <sup>[11]C</sup>SCH23390 | *kinfitr* | SRTM | 1.49 | 11.1 | 0.83 | 4.6 | 5.0 |
|  | PMOD | SRTM | 1.49 | 11.2 | 0.83 | 4.6 | 5.0 |
|  | *kinfitr* | ref Logan | 1.48 | 11.3 | 0.82 | 4.8 | 5.3 |
|  | PMOD | ref Logan | 1.56 | 12.3 | 0.80 | 5.6 | 7.0 |
|  | *kinfitr* | MRTM2 | 1.49 | 11.1 | 0.83 | 4.6 | 4.9 |
|  | PMOD | MRTM2 | 1.51 | 11.3 | 0.82 | 4.9 | 5.4 |

**Figure 1:**
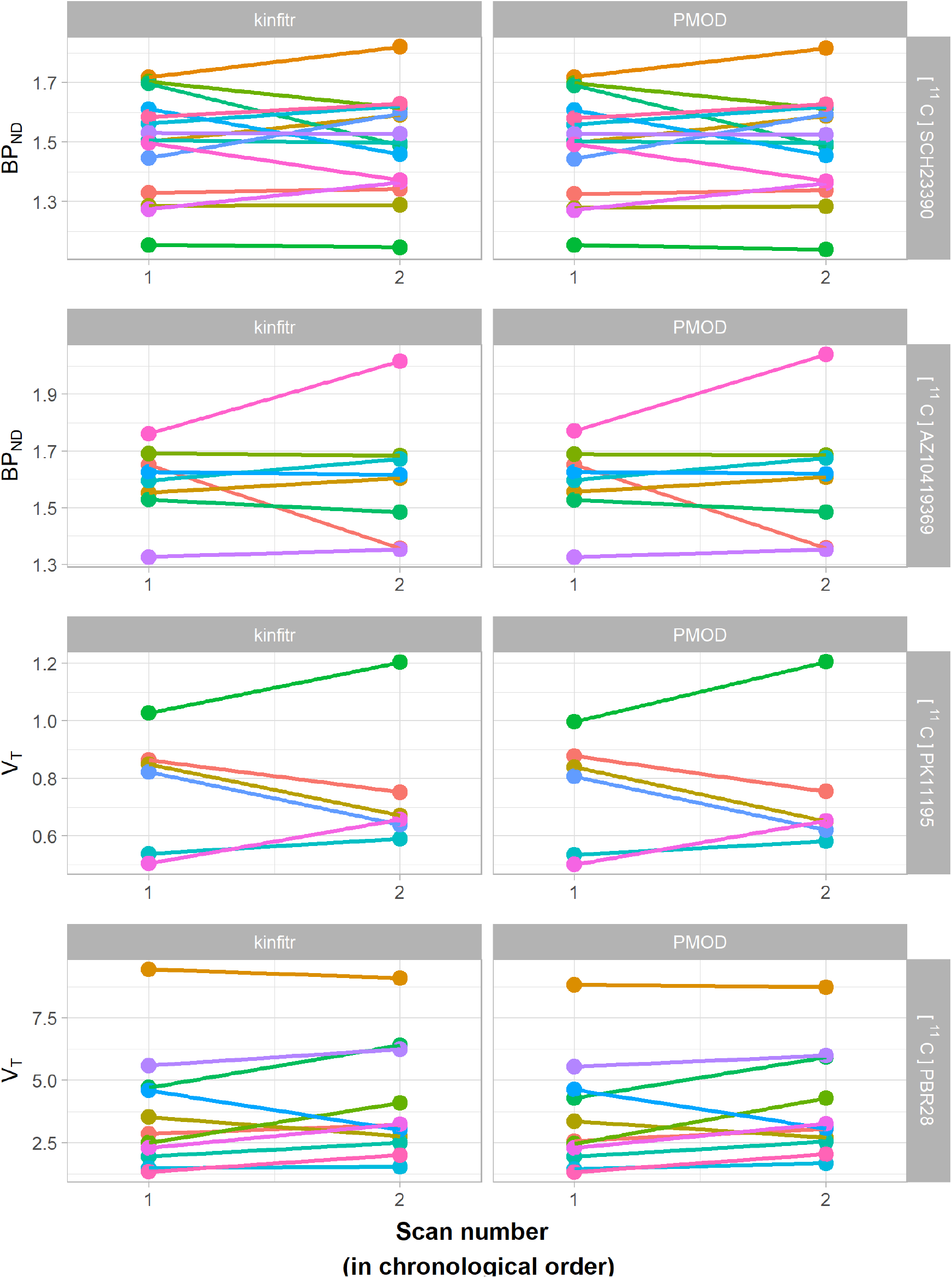
Binding estimate values comparing the first and second PET measurements. Each colour corresponds to a different individual, and the lines connect both of their two measurements. The ROI used in making this figure were the higher-binding regions for each radioligand in this study, i.e. the occipital cortex for [^11^C]AZ10419369, the striatum for [^11^C]SCH23390 and the thalamus for both TSPO ligands. The kinetic models represented here are SRTM for the estimation of BP_ND_ (above two rows), and the invasive model 2TCM for the estimation of V_T_.

### Microparameters

We also compared the values of microparameters (i.e. individual rate constants) estimated using the nonlinear methods. Figure 2 shows a comparison between the values of R1 and k_2_ obtained using SRTM for [^11^C]AZ10419369 and [^11^C]SCH23390. We observed Pearson’s correlation coefficients of >0.99 for both R1 and k_2_ estimated by *kinfitr* and PMOD. Similarly, the relationships between the microparameter estimates obtained using 2TCM for [^11^C]PBR28 and (R)- [^11^C]PK11195 were assessed (Figure 3). We found high correlations between *kinfitr* and PMOD estimates of K_1_, k_2_, k_3_ and k_4_ (mean Pearson’s correlation coefficients of >0.99, 0.81, 0.80 and 0.88 respectively).

**Figure 2:**
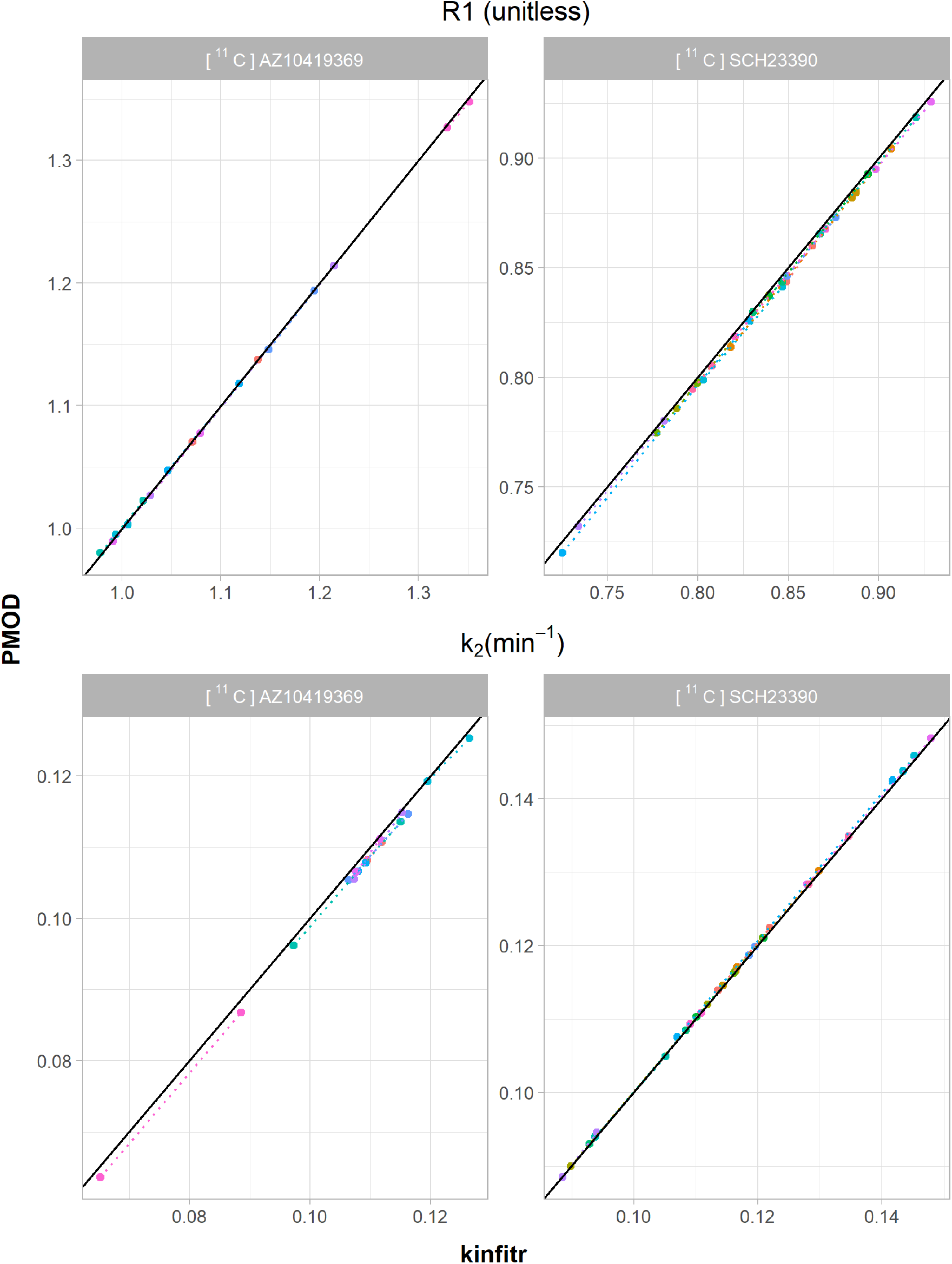
Microparameter comparison for the simplified reference tissue model (SRTM). The relationship between the values of individual rate constants calculated by either kinfitr or PMOD. The results for the radioligand [^11^C]AZ10419369 are derived from the occipital cortex ROI, whereas the results for [^11^C]SCH23390 correspond to the striatum. The diagonal line represents the line of identity. Each colour corresponds to a different subject, and the dotted lines connect both measurements from the same subject.

**Figure 3:**
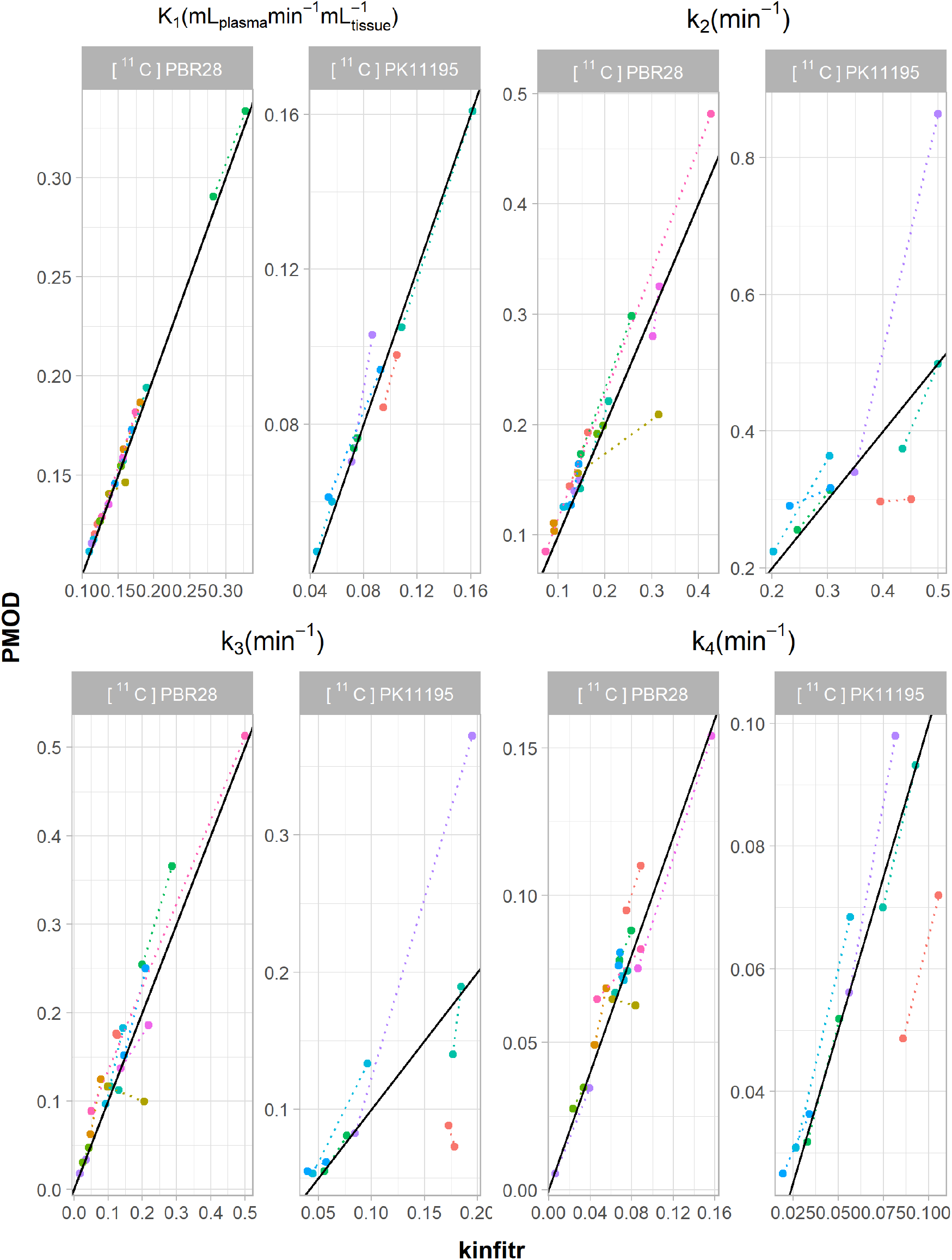
Microparameter comparison for the two-tissue compartment model (2TCM). The relationship between the values of individual rate constants calculated by either kinfitr or PMOD. All results were derived from the thalamus region. The diagonal line represents the line of identity. Each colour corresponds to a different subject, and the dotted lines connect both measurements from the same subject.

## Discussion

In this study, we evaluated the performance of *kinfitr* by comparing radioligand binding estimates to those obtained with the established commercial software PMOD. We assessed the similarity between these tools using four datasets, each encompassing a different radioligand, and employed three kinetic models for invasive and non-invasive applications. Mean regional BP_ND_ and V_T_ values computed by both tools were similar to those reported in previous literature on the same radioligands [32–34, 52]. We observed high agreement between estimates of BP_ND_ and V_T_ using *kinfitr* and PMOD. Furthermore, there were no substantial differences between the tools in terms of test-retest reliability for these measures. We further found that both tools exhibited a high degree of agreement in estimates of the microparameters, as well as high agreement between the estimates of the different models assessed using each tool separately. While the bias between some outcome measures estimated with the two tools was non-negligible (Table 1), the high correlations for all outcomes mean that this would not present an issue when using one or the other tool within a given dataset.

Despite the overall high similarity with regard to binding estimates, the linearized models (i.e. MA1, MRTM2 and both invasive and non-invasive Logan plots) exhibited a slightly lower degree of agreement the nonlinear models (2TCM and SRTM). This observation is most likely explained by the fact that the linearized models require the selection of a t* value, which was performed differently using the two tools. As described in more detail in the Supplementary Material S1, PMOD fits a t* value for the user, whereas *kinfitr* requires the user to specify a t* based on several plots as visual aids with which to select an appropriate value. As such, the PMOD interface makes it more convenient to fit t* values independently for each individual, while the *kinfitr* interface encourages selecting a t* value which is generally applicable across all study participants.

With regard to the user interface of the two tools, the most important difference is that *kinfitr* requires the user to interact with the data using code, while PMOD makes use of a graphical user interface (GUI), i.e. the user clicks buttons and selects items from drop-down menus. As such, *kinfitr* requires learning basic R programming before it can be used effectively, while PMOD can essentially be used immediately. Therefore, *kinfitr* may be perceived as having a steeper learning curve than PMOD. However, in our experience, *kinfitr* provides the user with greater efficiency once a moderate degree of proficiency has been gained. For instance, as a result of the code interface, re-running an analysis using *kinfitr* on all study participants using different parameters (e.g. altering a fixed v_B_ or t* value) or a different model, can be performed by modifying only the relevant lines of code. In contrast, performing re-analyses using PMOD can require a great deal of manual effort, as all tasks must essentially be repeated. This exemplifies the fundamental benefit of computational reproducibility: by crystallising all steps in computer code, the results can easily be generated anew from the raw input data. This has additional benefits, such as making the detection of potential errors substantially easier as all user actions are recorded transparently in the analysis code, and allowing others to more quickly and easily adapt, modify or build upon previous work.

It is important to note that the kinetic modelling was not performed in an identical manner between the two tools; rather we performed the modelling in a manner as consistent with the way users might actually use the software as possible. This was done in order to emphasize ecological validity. While this diminishes the extent to which we can specifically compare the outcomes using both of the two tools, our intention was instead to compare how both tools would be expected to perform independently in practice. This approach focuses on the extent to which outcomes might potentially differ between these tools, rather than the extent to which they can be made similar. It is thus reasonable to assume that even higher agreement could be achieved if additional measures were taken to make each analytic step identical.

In summary, we showed excellent agreement between the open source R package *kinfitr*, and the widely-used commercial application PMOD, which we have treated as a gold standard for the purpose of this analysis. We therefore conclude that *kinfitr* is a valid and reliable tool for kinetic modelling of PET data.

## Declarations

### Ethics approval and consent to participate

In all studies, the data collection was approved by the Regional Ethics and Radiation Safety Committee of the Karolinska Hospital, and all subjects had provided written informed consent prior to their participation.

### Consent for publication

Not applicable

### Competing interests

No conflict of interest relevant to the present work.

### Funding

S.C. was supported by the Swedish Research Council (Grant No. 523-2014-3467). J.T. was supported by the Swedish Society of Medicine (Svenska Läkaresällskapet).

### Authors’ contributions

GJM conceived of the study. LF was involved in planning and supervision of data collection. GJM and JT designed the study. JT and GJM analysed the data and interpreted the results. JT, GJM and SC drafted the article. All authors critically revised the article and approved of the final version for publication.

## Acknowledgements

The authors would like to express their gratitude to the members of the PET group at the Karolinska Institutet, for assistance over the course of the investigation, and all who participated in collecting the data which was used in the present study, in particular the first authors: Per Stenkrona, Magdalena Nord, Karin Collste and Aurelija Jučaite. In addition, we would like to thank Dr. Ryosuke Arakawa for his assistance in the use of PMOD and the interpretation of its manual.

## Notes

https://github.com/tjerkaskij/agreement_kinfitr_pmod

