## Supplementary Materials for "Kinfitr – an open source tool for reproducible PET modelling: validation and evaluation of test-retest reliability"

### Supplementary material

#### Supplementary Materials S1

#### Differences Between the analyses done in PMOD and *kinfitr*

#### PMOD analysis

<u>Interface and Auditability</u>

PMOD relies on a graphical user interface (GUI) (Figure 1). Data is sequentially loaded for each study participant, one-by-one, and the kinetic models are fitted using buttons and drop-down menus. User actions can be logged by purchasing the “audit trail license” (1).

<u>Weighting schemes</u>

We made use of the default weighting scheme of PMOD, which is constant weighting. This means that the same weight is applied to all data points. The whole brain TAC was used for the calculation of weights.

<u>t* values</u>

For the selection of a value for t*, PMOD allows the user to automatically select an appropriate value By default, the software will search for the earliest value of t* such that the deviation between the regression line and all of the observed values is less than 10% (2). The t* value was fitted using the higher-binding region for each tracer. This t* value was then used for the other regions for each tracer and model.

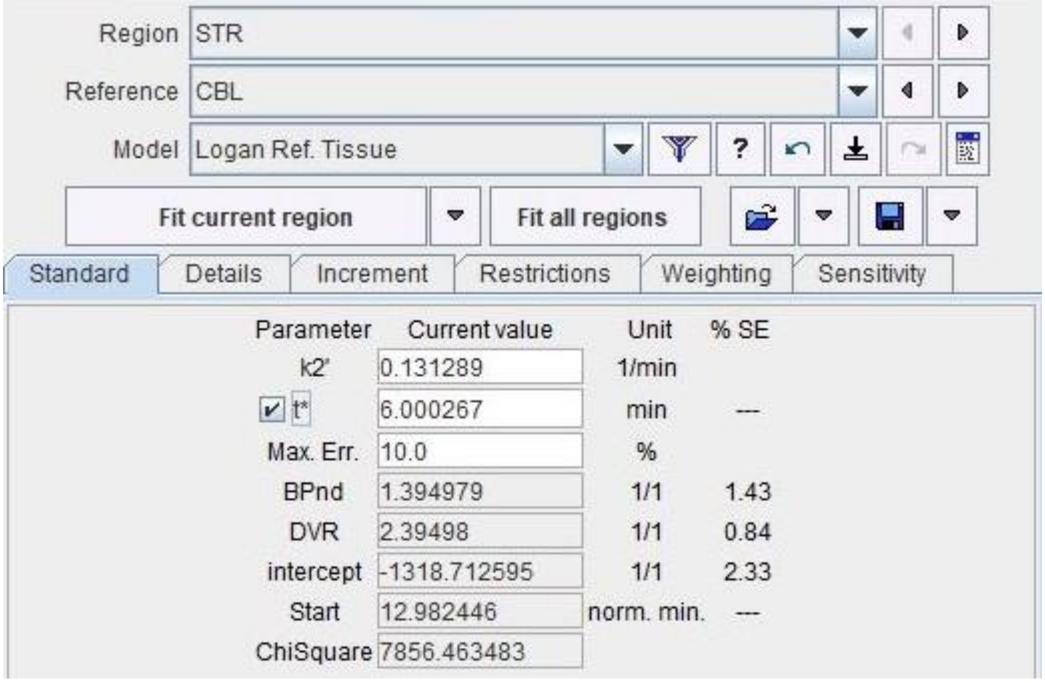


*This image shows the PMOD interface during the fitting of the reference tissue Logan plot for a study participant belonging to the $[^{11}C]$SCH23390 cohort. The t* button is checked, indicating that t* will be or has just been fit. The Max.Err criterion, set to 10%, indicates the cutoff value of maximum percentage residuals used to fit t*. The Striatum, being a high-binding region for $[^{11}C]$SCH23390, was used when fitting t* for all*

*study participants belonging to that cohort, and the cerebellum was used as a reference region. " k2' " was manually inserted after having estimated it using MRTM1.*

PMOD is therefore designed in such a way that fitting a t* value for each PET examination is made very convenient. If one were to wish to use a single t* value across multiple individuals based on the results from everyone, this would take more time and effort. One would sequentially load in the data for each subject individually, fit the model and t* value, and write down the different t* values obtained. Afterwards, one could evaluate these values and decide on an appropriate value for t*, and then sequentially load in the data for each subject once more, manually enter the t* value, and fit the model again using this value. For this reason, we suggest that the user interface encourages the use of individual t* values for each measurement, as this approach is much more convenient.

#### *kinfitr*

Interface and Auditability

In *kinfitr*, the user interacts with the software through R code. We made use of R notebooks, by which writing, code and code outputs are all interspersed with one another (3). These analysis notebooks are shared online (https://github.com/tjerkaskij/agreement_kinfitr_pmod). All user actions are therefore represented in the analysis code, and can be checked and audited directly. This is one of the central benefits of computational reproducibility.

One of the other proposed benefits of computational reproducibility is that code can be recycled and repurposed from project to project, or even between analysts. An example of this in practice is that the code within the analysis notebooks for [$^{11}$C]AZ10419369 and [$^{11}$C]SCH23390 is nearly identical and could therefore be copied from one to the other analysis and modified as necessary.

Weighting schemes

We used the default weighting scheme from *kinfitr*. This method defines weights as the square root of the product of the frame durations and the non-decay-corrected time activity curve of a large region. The resulting values are then taken as a proportion of the range between 0.7 and 1 to restrict their range. The whole brain TAC was used for the calculation of weights.

t* Values

In contrast to PMOD, selection of t* is not automated in *kinfitr*. Rather, the user is presented with $R^2$ values, maximum percentage residuals and changes in binding estimates for each potential value of t*, for three different regions of interest for each PET measurement, recommended to consist of regions with high, medium and low binding (Figure 2). In this way, the user must select an appropriate value for t* by examining all of the available information. It is further explicitly recommended in the documentation that these visual aids should be examined for several individuals/measurements to select a single t* value for the entire cohort.

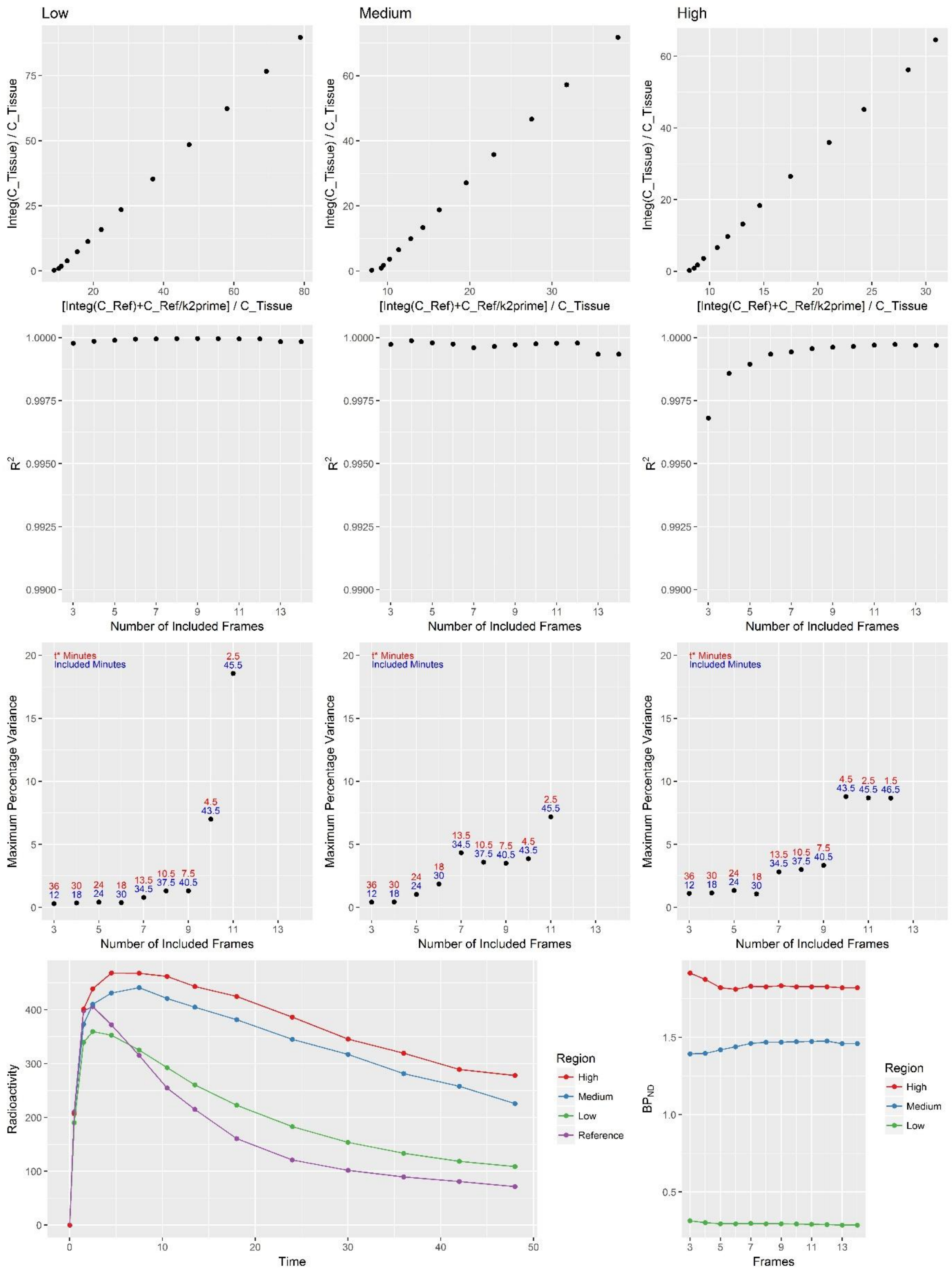


*Graphical aids provided by kinfitr for the selection of an appropriate t* value.*

In *kinfitr*, all data is loaded at once, and so producing all t* selection figures for several or all measurements in a cohort is straightforward. As such, *kinfitr* makes it more convenient to examine these t* visual aids, and to make a judgement about the most appropriate value for a t*, across the whole cohort. If, instead, one were to wish to select an individual value for t* for each individual, this would take a great deal more time and effort. One would have to examine the t* aids one-by-one, and write down the selected t* value for each, add the selected t* values to the data, and then re-run the modelling by pointing the model to the newly created t* column for which value to use.  Instead, this was a deliberate and opinionated design decision to avoid the potential for overfitting the selection of t* values, and to present the user with as much information as possible to make as informed a choice as possible.  For this reason, the user interface encourages the use of a single t* values for a whole cohort, as this approach is much more convenient, and is furthermore explicitly encouraged.

### Supplementary Materials S2

#### Demonstrating the outlier for PBR28 VT relative to the remainder of the data

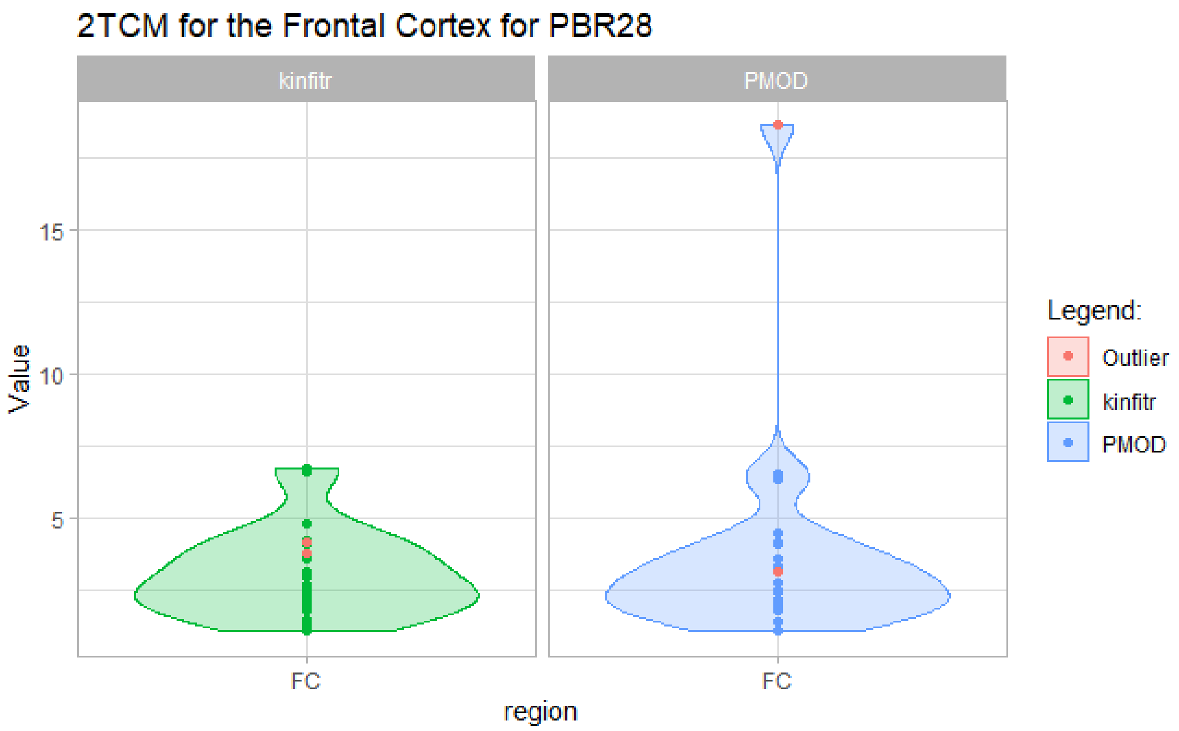


*Violin plot showing the results for 2TCM for the frontal cortex using the radioligand PBR28. The excluded study participant is marked in red. Results are shown for both kinetic modelling tools.*

### Supplementary Materials S3

#### Relationship between Binding Outcomes Estimated Using Each Tool

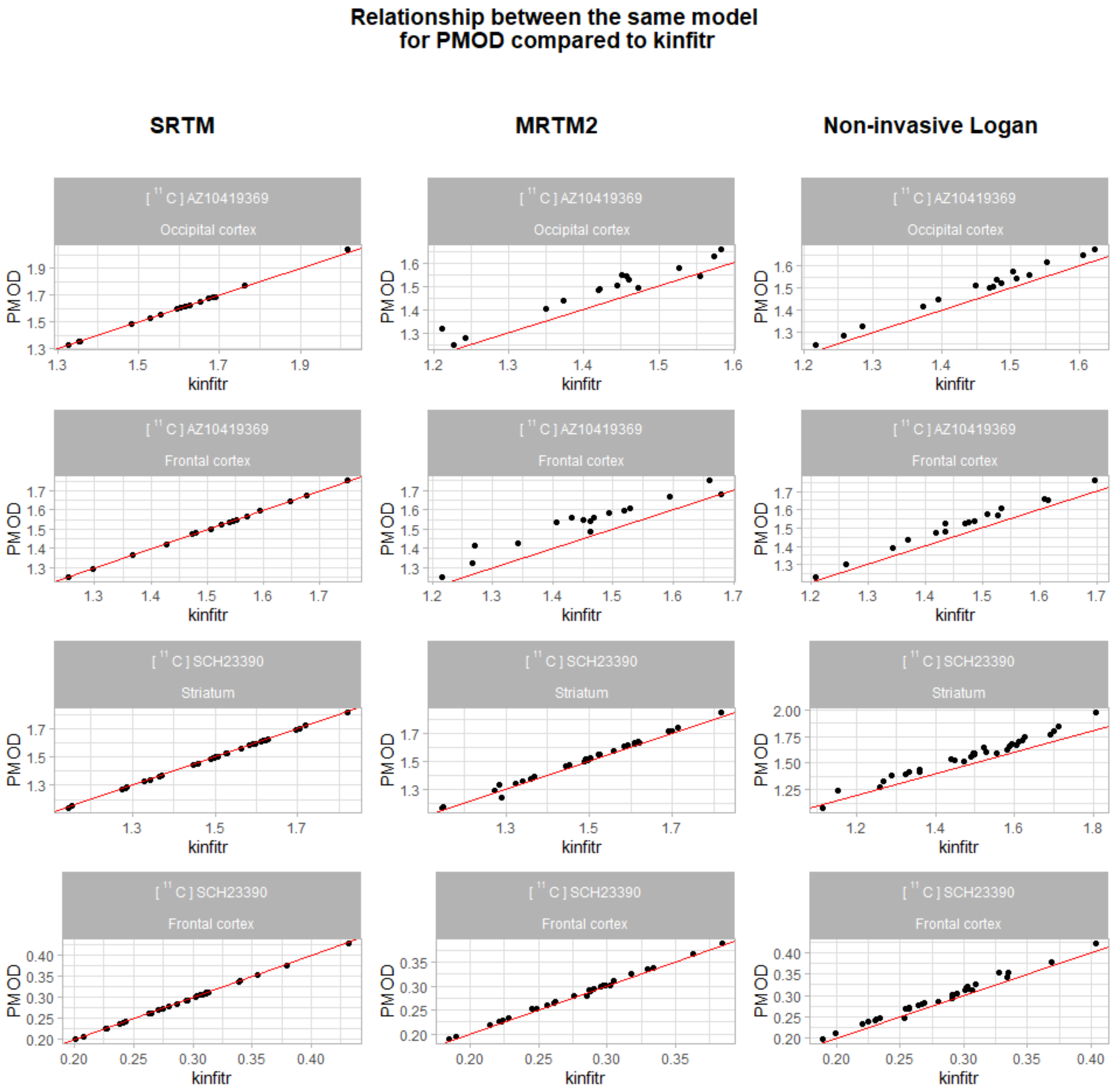


*This figure shows the relationship between the same model using kinfitr and PMOD for the radioligands SCH23390 and AZ10419369. The red diagonal line is the "line of identity", while the points represent the study participants.*

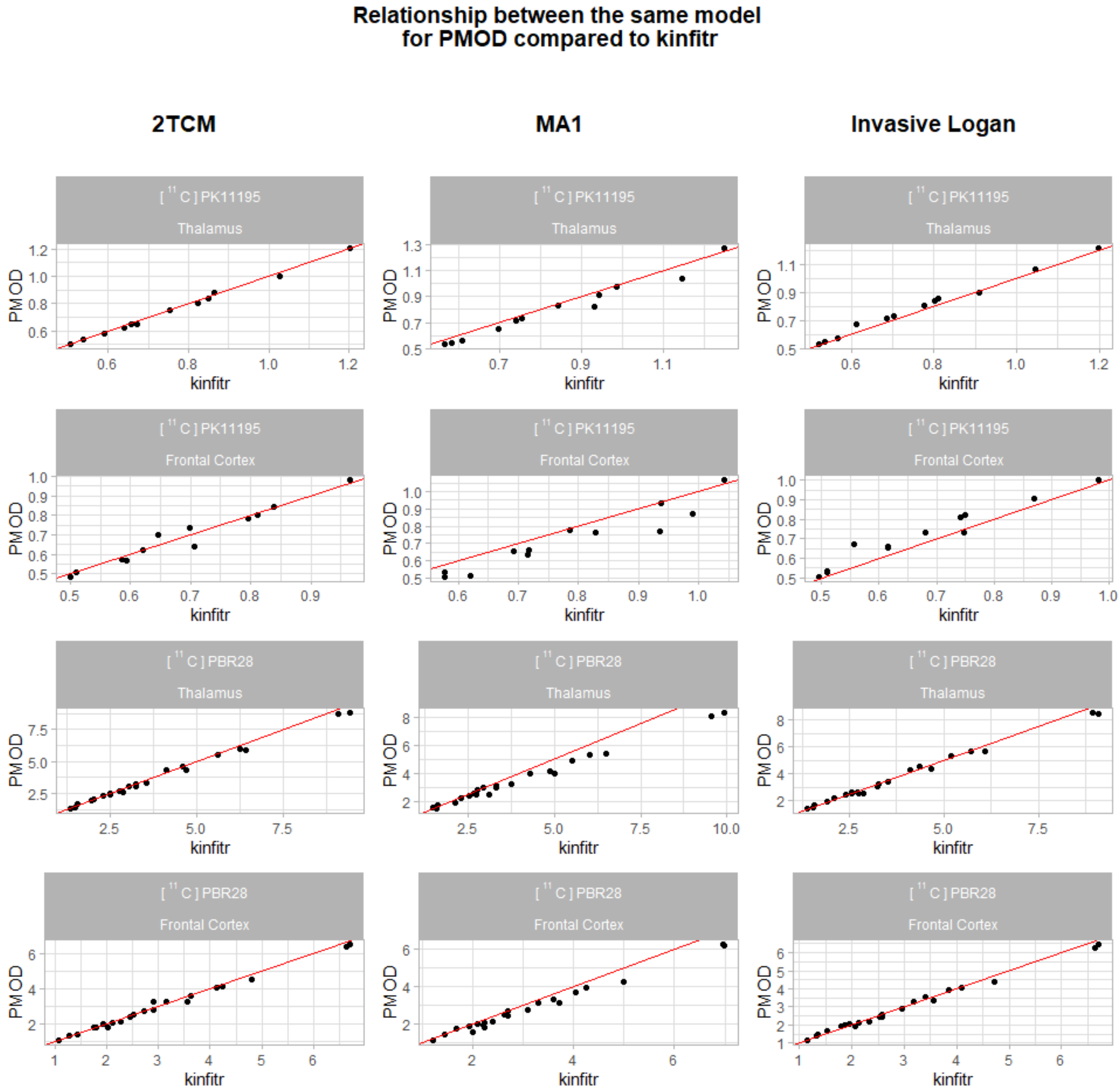


*This figure shows the relationship between the same model using kinfitr and PMOD for the radioligands PK11195 and PBR28. The red diagonal line is the "line of identity" while the points represent the study participants.*

### Supplementary Materials S4

#### Relationship between Binding Outcomes between Models Estimated Using Each Tool

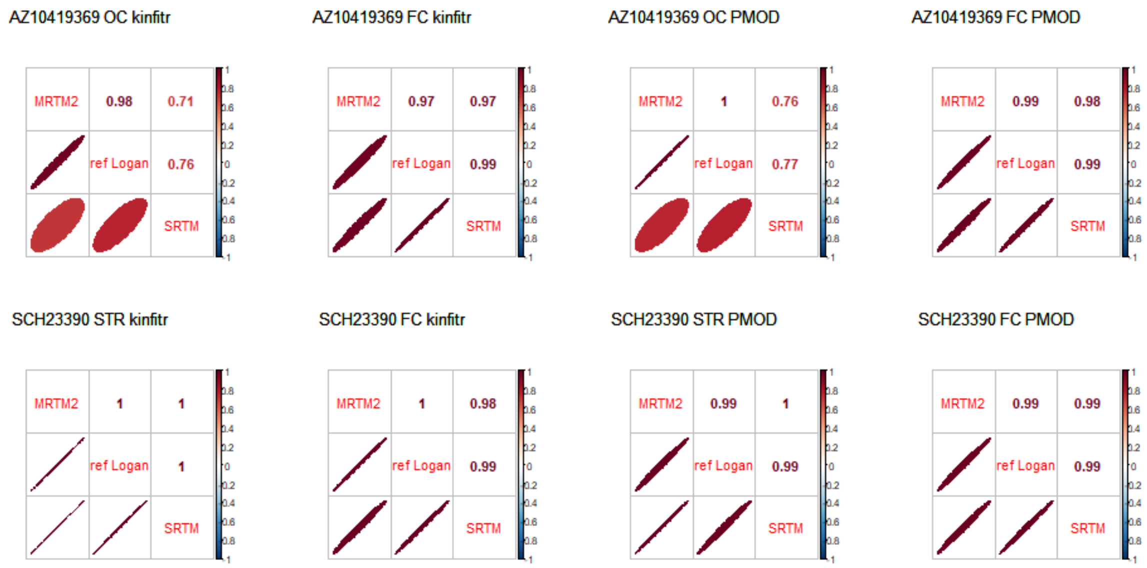


*Correlations between the three non-invasive kinetic models for each region and radioligand for both kinfitr and PMOD. Abbreviations: OC = Occipital cortex, FC = Frontal cortex, STR = Striatum.*

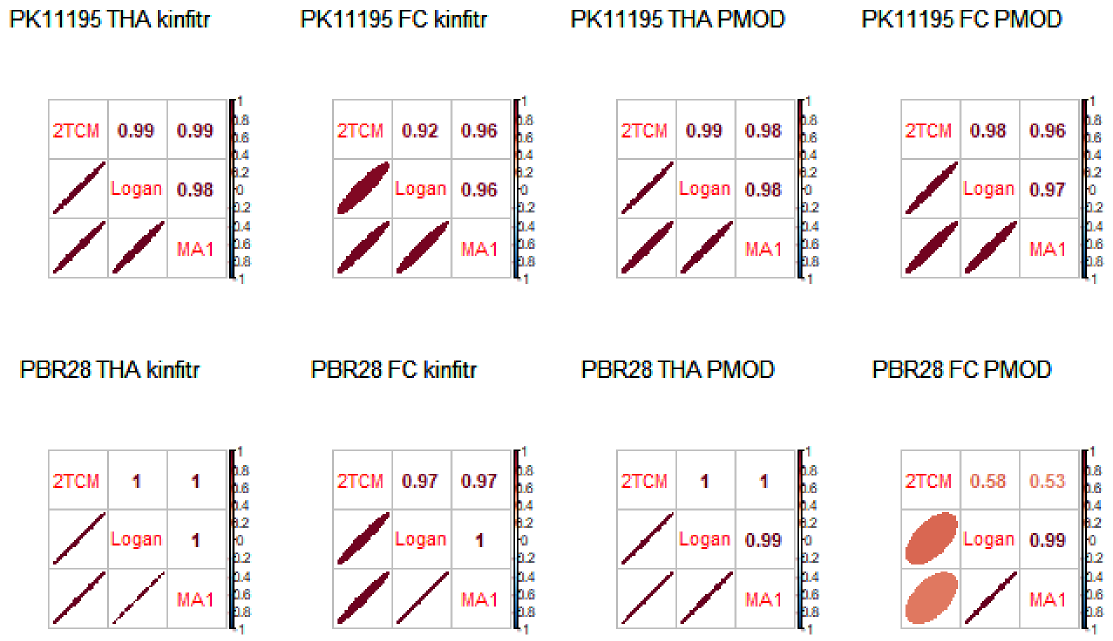


*Correlations between the three non-invasive kinetic models for each region and radioligand for both kinfitr and PMOD. Abbreviations: THA = Thalamus, FC = Frontal cortex.*

3. Xie Y (Mathematician), Allaire JJ, Grolemund G. R Markdown : the definitive guide. 303 p.
